# Estrogen Deficiency alters Vascularization and Mineralization dynamics: insight from a novel 3D Humanized and Vascularized Bone Organoid Model

**DOI:** 10.1101/2024.10.09.614903

**Authors:** Muhammad M.M. Bukhari, Mostafa Khabooshani, Syeda M. Naqvi, Laoise M. McNamara

## Abstract

Osteoporosis is not merely a disease of bone loss but also involves changes in the mineral composition of the bone that remains. *In vitro* studies have investigated these changes and revealed that estrogen deficiency alters osteoblast mineral deposition, osteocyte mechanosensitivity and osteocyte regulation of osteoclastogenesis. During healthy bone development, vascular cells stimulate bone mineralization via endochondral ossification, but estrogen deficiency impairs vascularization. Yet, existing *in vitro* bone models overlook the role of vascular cells in osteoporosis pathology. Thus, here we (1) develop an advanced 3D vascularized, mineralized and humanized bone model following the endochondral ossification process, and (2) apply this model to mimic postmenopausal estrogen withdrawal and provide a mechanistic understanding of changes in vascularization and bone mineralization in estrogen deficiency. We confirmed the successful development of a vascularized and mineralized human bone model via endochondral ossification, which induced self-organization of vasculature, associated with hypertrophy (collagen X), and promoted mineralization. When the model was applied to study estrogen deficiency, we reported the development of distinct vessel-like structures (CD31+) in the postmenopausal 3D constructs. Moreover, during estrogen withdrawal vascularized bone demonstrated a significant increase in mineral deposition and apoptosis, which did not occur in non-vascularized bone. These findings reveal a potential mechanism for bone mineral heterogeneity in osteoporotic bone, whereby vascularized bone becomes highly mineralized whereas in non-vascularised regions this effect is not observed.

**New and Noteworthy:** Here we develop an *in vitro* 3D vascularized and humanized bone model following an endochondral ossification approach. We applied the model to recapitulate estrogen deficiency as representative of osteoporotic phenotype. The results of this study reveal that estrogen deficiency exacerbates formation of 3D vessel like structures in vascularized models and thereby drives mineral deposition.

## 1. Introduction

Osteoporosis is a debilitating bone disease, in which severe bone loss occurs leading to fractures of the hip, wrist or vertebrae. The disease occurs in approximately 30% of postmenopausal women, when the dramatic reduction in circulating estrogen causes an imbalance in bone remodeling activity, whereby bone resorption exceeds formation, leading to overall loss in bone volume [1]. Previous research has revealed that osteoporotic bone loss is also accompanied by fundamental changes in bone mineralization and cellular responses to mechanical loads but the mechanisms underpinning these changes are not yet fully understood [2–6].

Estrogen is an important regulator of bone physiology and all bone cells possess receptors for estrogen (ERα and ERβ), which regulate the metabolic activity of osteoblasts, osteoclasts and osteocytes [7]. Estrogen supplementation to bone cells enhances matrix synthesis [8], stimulates osteogenic signaling [9], increases osteoblast differentiation [10, 11] and inhibits osteoclastogenic activity [12–14]. Ovariectomy induced estrogen deficiency causes heterogeneity in bone mineral distribution [3], secondary mineralization [6] and also alters the mechanical environment of bone cells [15]. During recent years, studies have sought to unravel the molecular mechanisms underpinning these changes. *In vitro* 2D and 3D bone models of postmenopausal osteoporosis have been developed, where bone cells are accustomed to estrogen supplementation for a period of time, after which estrogen is withdrawn to recapitulate the postmenopausal condition [16–22]. Through this approach, it has been reported that estrogen withdrawal alters osteocyte morphology, mechanosensation [16], calcium signaling [17], induces osteocyte apoptosis [18] and osteocyte mediated osteoclastogenesis [19]. *In vitro* estrogen-deficient MLO-Y4 cells activated hedgehog and osteoclastogenic paracrine signaling and exhibited an increase in pro-osteoclastogenic gene expression (RANKL/OPG ratio) in response to fluid flow [20]. Even the conditioned media from mechanically stimulated and estrogen deficient osteocytes increased CTSK gene expression and TRAP activity of osteoclasts [22]. Estrogen deficiency causes an increase in the expression of SOST in osteocytes, which increases osteoclastogenesis by inhibiting the Wnt/β-catenin signaling pathway [22]. Recently, a 3D model of bone was developed by encapsulating murine osteoblasts and osteocytes in gelatin, and this model was applied to reveal that estrogen deficiency and mechanical loading stimulated osteocyte differentiation, matrix mineralization, and regulated osteoclastogenesis [21].

There have been significant developments in bioengineered 3D *in vitro* models of tissue for disease modelling and precision medicine. Murine cell lines have been used to develop models of osteocyte differentiation but have not shown to develop fully mineralized matrix [23]. *In vitro* bone models have encapsulated human bone marrow stem cells in 3D hydrogels (gelatin, alginate and fibrin) and that are differentiated to develop mineralized matrix through the use of osteogenic supplements (dexamethasone, ascorbic acid and βglycerol phosphate) in the culture media [24–27]. In a recent study, a 3D bone-like organoid was developed by seeding bone marrow stems cells on silk fibroin and culturing them in dynamic culture, which induced stem cell differentiation into osteoblasts and osteocyte-like cells, extracellular matrix synthesis, osteocyte network formation and communication via sclerostin expression [28]. Although these *in vitro* studies provide insights into the molecular mechanisms underpinning early stage osteogenesis, previous models do not mimic the complex multicellular and vascularized microenvironment of the bone.

Vascularization is precursor to endochondral bone formation, whereby vasculature invades the chondrogenic template during endochondral ossification [29], and initiates bone formation [30]. Vascular cells increase mineralization of chondrogenic template in 3D organoid models, which was shown to be related to hypertrophic chondrocyte release of collagen X and VEGF that drive the vascularization of the chondrogenic template and enhance mineralization [31, 32]. Vascular cells also possess receptors for estrogen [33], and a recent study revealed that ovariectomy induced estrogen deficiency reduces the number of type H blood vessels and bone volume in mice femur [34]. However, existing *in vitro* bone models for studying osteoporosis overlook the role of vascular cells in osteoporosis pathology.

The objectives of this study were to (1) develop an advanced 3D vascularized, mineralized and humanized bone model, following the endochondral ossification process to vascularize the chondrogenic template and drive mineralization, and (2) apply this model to provide a mechanistic understanding of changes in vascularization and bone mineralization during estrogen deficiency, representative of the postmenopausal condition.

## 2. Methods

The research builds upon our previous studies that implemented endochondral priming to promote mineralization in 3D organoid models [31, 32]. Here, we develop an advanced 3D vascularized and humanized bone model, combining human cell culture and following the endochondral ossification process to drive mineralization.

### 2.1. Cell culture

Human bone marrow mesenchymal stem cells (HBMSCs) were isolated from bone marrow (42 and 22-year-old females) under ethical approval and informed consent (Research Ethics Committee, University of Galway). The HBMSCs were expanded in minimum essential medium alpha (αMEM D4526; Merck, Germany) supplemented with 10 % fetal bovine serum (FBS; F7524; Merck, Germany) and 2% Antibiotic Antimycotic Solution (AAS, A5955; Merck, Germany), 2% L-glutamine (L-Glutamine, G7513; Merck, Germany) at 37 °C and 5 % CO_2_. The cells were trypsinized at 90% confluency by using 0.25 % trypsin ethylenediaminetetraacetic acid (EDTA; Invitrogen). HBMSCs were further cultured to passage 3–5, before using them in the experiment. HUVECs were commercially available and purchased from Lonza (Maryland, USA) and cultured in Clonetics endothelial cell basal medium (EGM2, C-2221; Promocell, USA). Upon reaching 80–90 % confluency, cells were passaged using 0.25 % trypsin ethylenediaminetetraacetic acid (EDTA; Invitrogen). HUVECs were cultured to passage 4, before using them in experiments.

### 2.2. Development of advanced 3D humanized and vascularized models of healthy and osteoporotic bone

We have previously established gelatin-based 3D cell culture systems and optimized the cell seeding density (2×10^6^ cells/ml), stem and substrate stiffness (0.58 kPa) for osteogenic differentiation [35, 36] and also the stem cell and endothelial cell ratios in co-culture to stimulate proliferation, survival, osteogenesis and angiogenesis [31]. Building on these studies, here HBMSCs (P5) were suspended in sterile microbial transglutaminase (mtgase; Activa WM, Aginomoto foods, Japan). The mtgase/cell suspension was used to crosslink an equal volume of 6% gelatin (type A, 175 Bloom) (ratio of 1:1). The final concentration of mtgase and gelatin in the suspension, was 3% v/v. To make gelatin based 3D cellular constructs, 120µL of the gelatin-mtgase cell suspension was pipetted into custom made polydimethylsiloxane (PDMS) rectangular wells and allowed to cool at 4 °C for 8-12 minutes. These constructs were primed for 21 days (until day −21) with chondrogenic differentiation media (Dulbecco’s modified Eagle’s medium (DMEM) (Merck), 10 ng/mL transforming growth factor (TGF)-β3 (Invitrogen), 50 μg/mL ascorbic acid (Merck), 4.7 μg/mL linoleic acid-oleic acid (Merck), 100 nM dexamethasone (Merck) and 1× insulin–transferrin–selenium (ITS; Invitrogen)) (Fig. 1). At day −21 the 3D vascular constructs were developed by culturing the chondrogenic template within a layer (120µL) of gelatin-mtgase in which HUVECs (P4) and HBMSCs (P5) were encapsulated at 1:1. Non-vascularized constructs acted as controls and were developed by culturing the chondrogenic template within a layer of (120 µL) of gelatin-mtgase in which only HBMSCs were added. The cell seeding density in the final constructs was 2 × 10^6^ cells/ml (240,000 cells/construct).

**Figure 1:**
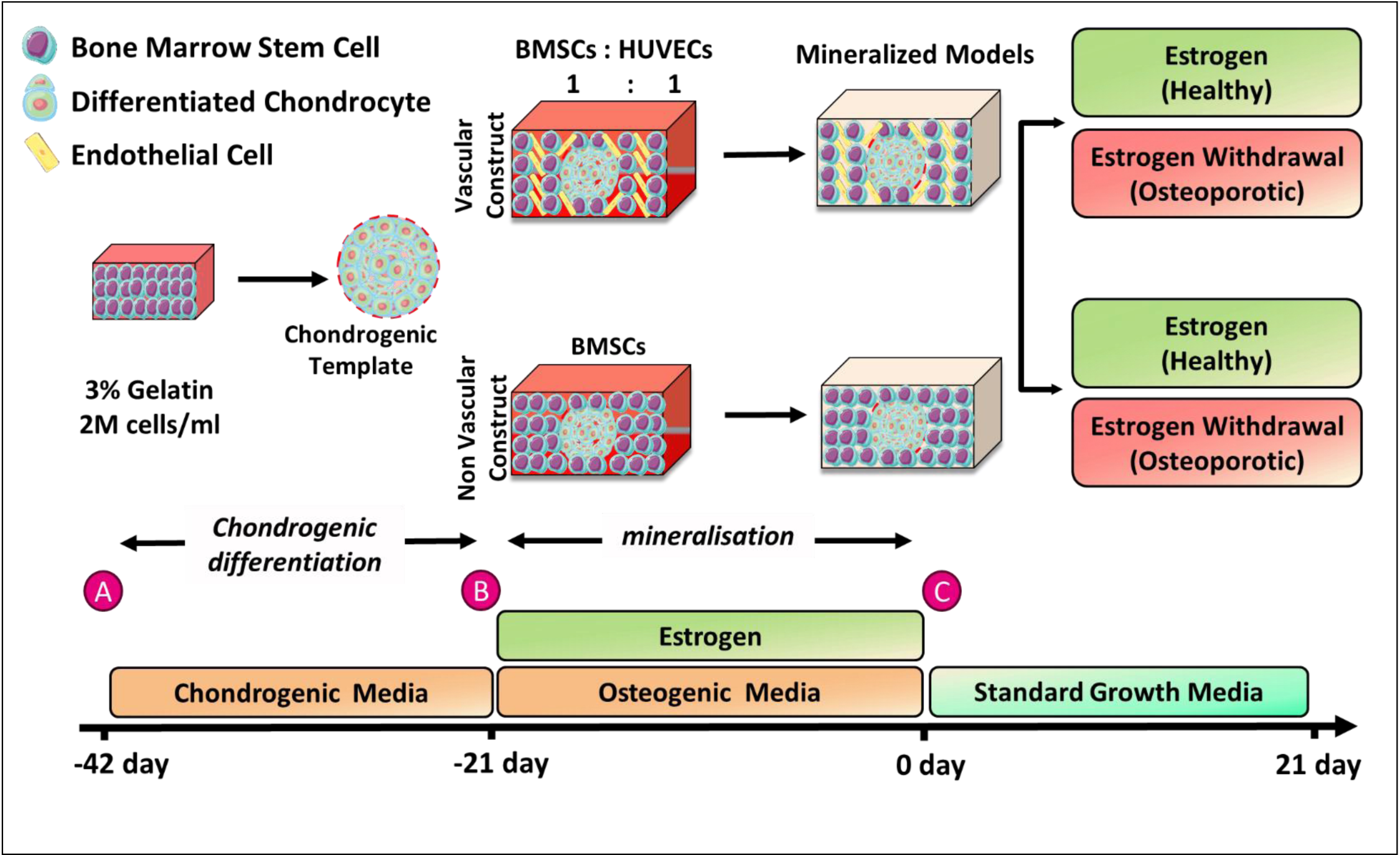
Experimental design. **(a)** Development of chondrogenic template by differentiating HBMSCs in gelatin-mtgase constructs (from −42 day to −21 day), **(b)** Culture of chondrogenic template in vascular (HBMSCs: HUVECs) and non-vascular (BMSCs) gelatin-mtgase constructs under osteogenic differentiation and estrogen supplementation (from −21 day to 0 day), **(c)** Mineralized vascular and non-vascular models at day 0 (day 43 in culture)and further culture under estrogen supplementation or estrogen withdrawal (from 0 day to 21 day).

Next, the vascular and non-vascular constructs were cultured in osteogenic media (EGM2 media (Promocell) for a further 21 days until day 0 (day 43 in culture), supplemented with 10nM β-estradiol, 10% FBS, 100 nM of dexamethasone, 50 mg/mL ascorbic acid, and 10 mM βglycerol phosphate (Merck). At day 0 (day 43 in culture) osteogenic media was stopped and thereafter vascular and non-vascular constructs were cultured for further 21 days in EGM2 media (Promocell) with either continued estrogen or without estrogen. The samples were collected and analyzed at day 0, day 10 and day 21 (Fig. 1).

### 2.3. Sample collection and storage

The 3D constructs were analyzed using biochemical assay, histochemical staining, immunostaining and RT-PCR, at day −21, 0 (day 43 in culture), 10 and 21. The culture medium from the 3D constructs was collected, frozen, and stored at −80°C until biochemical assays could be performed. The construct for the DNA assay, calcium assay and RT-PCR were washed with phosphate-buffered saline (PBS) and were frozen at −80°C until analyzed. The constructs for the immunofluorescence and histochemical analysis were washed with 1X PBS and fixed in paraformaldehyde overnight and stored in 1X PBS, at 4°C until analyzed. Three independent experiments were carried out with at least two replicates in each experiment (n; biological replicate, n = 3 for histological and immunofluorescence analysis, n = 6 for calcium assay, n = 6 for DNA assay, n=6 for ALP assay and n=5 for rtPCR).

For immunofluorescence staining, the cells within the constructs were permeabilized with 0.5% Triton-X in PBS for 35 minutes at 4°C under agitation. After three washing steps in 1% BSA, the constructs were immersed in 1% BSA blocking solution for 1 hour under agitation. The respective constructs were incubated with primary antibody (see table 1) overnight. Next day, the constructs were washed with 1%BSA three times with 10 minutes’ incubation during each wash. After that, the constructs were incubated for 1 hour under rotation with fluorescent secondary antibodies respective to the primary antibody used (description in table 1). The constructs were again washed with 1% BSA to remove the unbound secondary antibodies. The constructs were counter stained with phalloidin-TRITC (1:500) to stain the actin cytoskeleton and DAPI (1:2000) to stain the nuclei. Negative controls (no primary antibody controls) for sclerostin and DMP-1 staining were also prepared by omitting the primary antibody during the staining process. The stained 3D constructs were imaged by using Fluoview 3000 (FV3000) confocal laser scanning microscope system (Olympus) at either 20X (oil) or 60X (oil) lenses. The Z-stack images were captured with step size 1 µm or 5 µm. NIH ImageJ software was used to analyze the images.

**Table 1.** List of primary and secondary antibodies.

|  | Primary Antibody | Dilution | Secondary Antibody | Dilution |
| --- | --- | --- | --- | --- |
| 1 | Ab-6308 Mouse monoclonal [COL-1] | 1:250 | Ab-7067 goat anti mouse IgG (Biotin conjugated) | 1:1000 |
| 2 | Ab-34712 Rabbit polyclonal [COL-2] | 1:200 | Ab-207995 goat anti rabbit IgG (Biotin conjugated) | 1:500 |
| 3 | Ab-49945 Mouse monoclonal [COL-X] | 1:500 | Ab-5929 Goat anti-mouse IgM (Biotin conjugated) | 1:500 |
| 4 | Ab-85799 Rabbit polyclonal [Sclerostin] | 1:250 | Dylight 488 goat anti-Rabbit (Jackson Immunoresearch) | 1:500 |
| 5 | Ab-106100 Rat monoclonal [Endomucin] | 1:50 | Ab-150155 Donkey anti rat [Alex Fluor 647] | 1:100 |
| 6 | Ab-28364 Rabbit polyclonal [CD31] | 1:50 | Dylight 488 goat anti-Rabbit (Jackson Immunoresearch) | 1:100 |
| 7 | Sc-73633 Mouse monoclonal [DMP-1] | 1:250 | A-11005 Goat anti-Mouse IgG (H+L), Alexa Fluor™ 594 | 1:500 |
| Table 1. List of primary and secondary antibodies. |  |  |  |  |

### 2.4. Confirmation of cell numbers and vascular development

The total DNA was quantified using a colorimetric assay. Briefly, the constructs were digested with 3.88 U/mL papain in 0.1 M sodium acetate, 5 mM L-cysteine-HCl, 0.05 M EDTA, pH 6.0 at 60°C under constant rotation for 18 h. Cell number was evaluated using the Hoechst 33258 DNA assay as previously described [37].

The immunostaining protocol was followed as described earlier, after blocking with bovine serum albumin, the constructs were incubated with the rabbit polyclonal anti-CD31 antibody polyclonal (ab28364, 1:50) and rat monoclonal anti-endomucin (ab-106100, 1:50) overnight. After that, the constructs were incubated for 1 hour under rotation with fluorescent secondary antibodies respective to the primary antibody used (description in table 1). The constructs were again washed with 1% BSA to remove the unbound secondary antibodies. The constructs were counter stained with DAPI (1:2000) to stain the nuclei. “No primary antibody” controls for CD31 and EMCN staining were also prepared by omitting the primary antibody during the staining process. The stained 3D constructs were imaged by using Fluoview 3000 (FV3000) confocal laser scanning microscope system (Olympus) at either 20X (oil) or 60X (oil) lenses. The Z-stack images were captured with step size 1 µm or 5 µm. NIH ImageJ software was used to analyze the images.

### 2.5. Confirmation of osteogenic differentiation and mineralization

#### 2.5.1 ALP production

Extracellular ALP production was quantified using a colorimetric assay of enzyme activity (SIGMAFAST p-NPP Kit; Merck), which uses p-nitrophenyl phosphate (pNPP) as a phosphatase substrate with ALP enzyme (Merck) as a standard. 50 µL pNPP solution and 40µL sample medium was added into a 96-well plate in triplicate. 10µL ALP enzyme was added into the standard wells. The plate was shielded from direct light by using aluminum foil and kept at room temperature for 1hour. The plate was read at 405 nm in a microplate reader.

#### 2.5.2 Calcium content

Total calcium deposition in the 3D constructs was quantified by using the Calcium Liquicolor kit (Stanbio Laboratories) according to the manufacturer’s protocol. The 3D constructs were digested by adding 1 mL of 1 M HCL and incubating at 60°C under rotation 10 rpm for 16 hours. The calcium standards were prepared according to the manufacturers protocol. Next, 10 µL of the digested samples was added to a 96-well plate in triplicates. Then 200 µL of the working solution was added into each well. The plate was shielded from the direct light by using aluminum foil and incubated at room temperature for 10 minutes. The plate was analyzed on a microplate reader at an absorbance of 550 nm.

#### 2.5.3 Histology

The constructs for the histological analysis were fixed in 4% paraformaldehyde overnight. Cryosectioning of the constructs was performed by embedding samples in OCT solution. Briefly, the constructs were incubated in the 5% sucrose overnight at 4°C. Next day, samples were incubated in 20% sucrose for 6 hours. After that the constructs were properly oriented into the molds and optimal cutting temperature compound (OCT) was added into the molds and kept at 4 °C for 20 minutes to allow the compound infiltration into the constructs. After 20 minutes the OCT embedded construct molds were stored in the −80 °C until sectioned. The constructs were sectioned with a thickness of 12µm by using Cryostat machine (Leica). The sections were stained with Alcian blue for sGAG, H & E for construct morphology, immunohistochemistry for collagen (II, I, X) and Von kossa staining for mineral.

#### 2.5.4 Immunohistochemistry

The sections were analyzed by immuhistochemistry staining for collagen II, collagen I and collagen X. The slides were rinsed with PBS and treated with chondroitinase ABC for 1 hours in humid box. Then the slides were rinsed with PBS and non-specific binding was blocked with 1% bovine serum albumin. Sections were incubated overnight in a humid box at 4°C with the primary antibody; mouse monoclonal anti-collagen I antibody (1: 250) or rabbit polyclonal anti-collagen type II (1: 250) or mouse monoclonal anti-collagen X antibody (1: 500). Next day, the slides were rinsed with PBS and the peroxidase activity of the sections was stopped by incubating with hydrogen peroxide. The sections were incubated for 1 h in the secondary antibody respective to the primary antibody used (description in the table 1). Color was developed using the Vectastain ABC reagent followed by exposure to peroxidase DAB substrate kit. Positive and negative controls of human ligament and cartilage were included for each antibody (supplementary fig 2)

#### 2.5.5 Polymerase Chain Reaction (PCR)

Total RNA was isolated with a custom CTAB method (Koster et al., 2016). Briefly, each construct was digested with 250 µL of RNAse-free proteinase K solution 5 mg/mL, containing 5 mM CaCl_2_. The digested constructs were mixed with 500 µL of CTAB solution (2% [w/v] Cetyl trimethylammonium bromide, 1.4 M NaCl, 20 mM EDTA, and 100 mM Trims, pH 8) containing 1% (v/v) β-mercaptoethanol. Then 500 µL of chloroform was added to each construct and centrifuged at 14,000 g for 2 min at room temperature. The upper/aqueous phase was transferred to a fresh tube and 800 µL of isopropanol (Fisher) was added to precipitate total RNA by centrifugation (14,000 g, 15 min, room temperature). The pellet was washed with 600 µL 70% (v/v) ethanol and dissolved in 20 µL low TE buffer (Thermo Fisher Scientific) for 15 min at 65°C. RNA purity and yield were assessed using a spectrophotometer (DS-11 FX, DeNovix), with 260/280 and 260/230 ratios over 1.8 for all constructs. 1 µg of RNA from each construct was transcribed into cDNA using a cDNA synthesis kit (Thermofisher, K1622) and thermal cycler (5PRIMEG/O2, Prime). Conventional PCR and agarose gel electrophoresis was conducted to confirm the specificity and annealing temperatures of the primers. The relative gene expression was studied by semi quantitative real-time polymerase chain reaction (sqRT-PCR, QuantStudio5) and Maxima SYBR Green/ROX qPCR Master Mix (2X) (Thermofisher, K0221).

### 2.6. Statistical Analysis

Statistical analyses were performed using GraphPad Prism (version 5) software. Two−way ANOVA was used for analysis of variance with Bonferroni’s post hoc tests to compare between groups. The results are presented as box plots with median, upper quartile, lower quartile, and interquartile range with all the data points visible. Significance was accepted at a level of p ≤ 0.05. The entire experiment was repeated three times.

## 3. Results

### 3.1. Development of a healthy multicellular, vascularized and mineralized human bone model

In the present research, we replicated the key events of the endochondral ossification process to develop a humanized, vascularized and mineralized bone mimetic model (Figure 2). Firstly, we developed the chondrogenic template by differentiating HBMSCs in 3D gelatin constructs. Chondrogenesis was confirmed by alcian blue staining at day 21, which showed a higher amount of glycosaminoglycans in the chondrogenic constructs compared to the non-chondrogenic groups (Supplementary Fig. 1). Subsequently, these templates were cultured within vascularly-primed gelatin layers for a further 42 days to drive vascularization and mineralization. The vascular layers developed multicellular connections with the chondrogenic template, which were visible with H&E staining as multicellular connections between the outer gel and the chondrogenic template (Fig. 2 b). Immunofluorescence staining for CD31 and EMCN confirmed the presence of type H (CD31^+^Emcn^+^) endothelial cells in the 3D constructs (Fig 2 c, d). Immunohistochemistry staining confirmed the deposition of collagenous proteins (Col-II, Col-X, Col-I) in the chondrogenic template characteristic of early-stage endochondral ossification (Fig 2 e, f, g). The constructs also stained positive for non-collagenous proteins, an osteoblast differentiation marker, dentin matrix acidic phosphoprotein I (DMP-I), and the osteocyte marker sclerostin (Fig 2 h, i). Von Kossa staining revealed mineralized nodules in the center of the chondrogenic template in vascular constructs (Fig 2 j).

**Figure 2:**
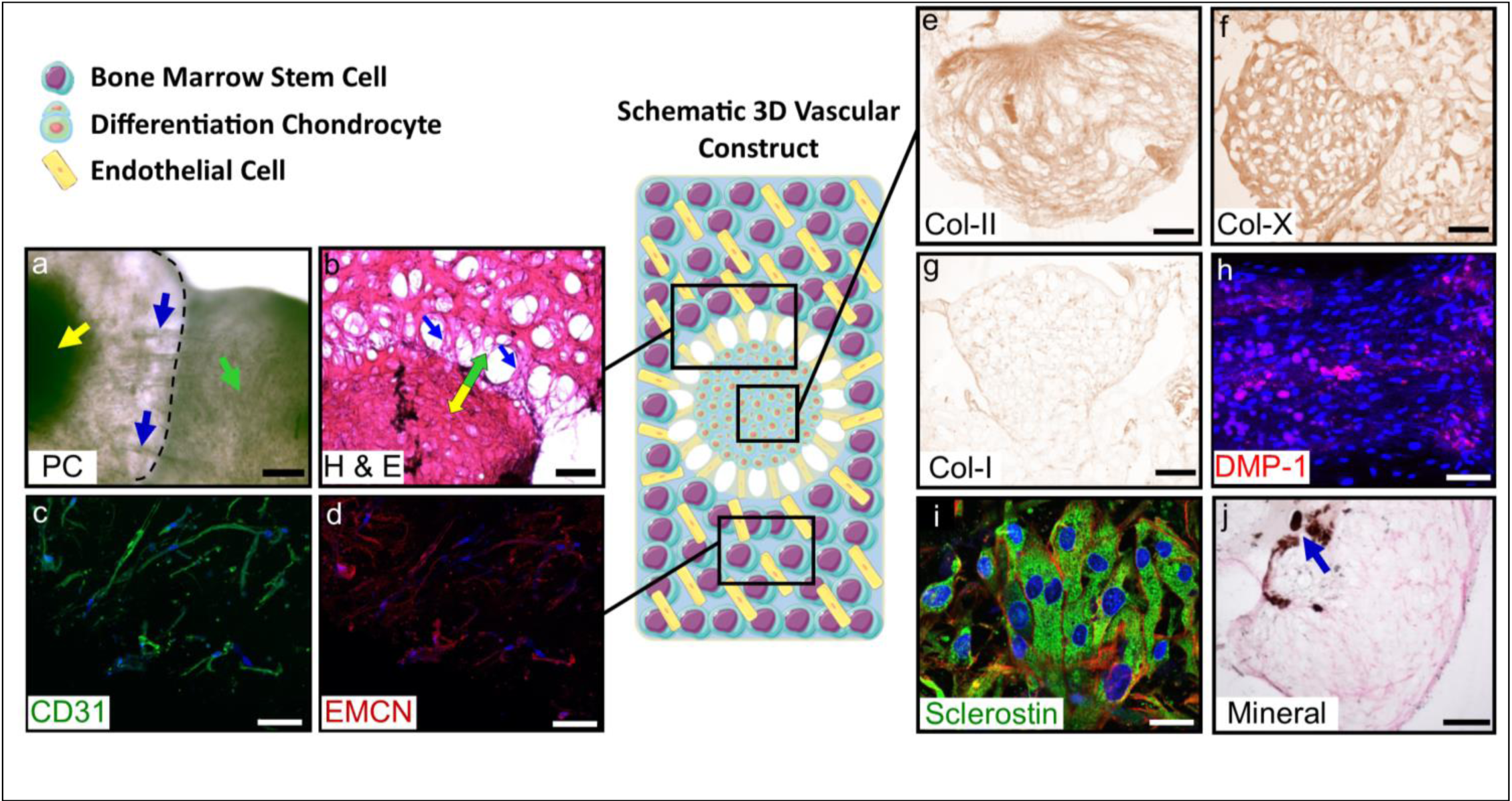
Development of a healthy (estrogen supplemented) vascularized and mineralized human bone model via endochondral ossification. **(a, b)** H&E staining at day 0 (day 43 in culture), chondrogenic template (yellow arrow) confirms integration of the vascular primed gelatin layer (green arrow) with the chondrogenic template through multicellular connections (blue arrows) (scale bar 500 µm). By day 21 immunofluorescence and immunohistochemistry staining confirmed **(c, d)** the retention and distribution of CD31^+^Emcn^+^ type H endothelial cells: CD31 (green), endomucin (red), DAPI (blue) (scale bar 100 µm) at day 21, **(e, f, g)** extracellular matrix production: collagen II, X and I (scale bar 500 µm), **(h, i)** osteoblast differentiation towards osteocytes: DMP-I (red), DAPI (blue) (scale bar 100 µm), sclerostin (green), actin (red), DAPI (blue) (scale bar 30 µm), **(j)** and mineral deposition: Von kossa staining for mineral, mineral nodule (blue arrow) (scale bar 500 µm).

### 3.2. Vascular priming under estrogen supplementation promotes osteocyte differentiation and mineralization, and delays hypertrophy and apoptosis

Our results uncover the influence of vascular priming under estrogen supplementation, which were reported to promote osteogenic differentiation and mineralization. Specifically, at days 10 and 21 all the groups under continued estrogen supplementation (healthy condition) stained positive collagen I, DMP-I, sclerostin and Von Kossa (mineral staining), see Figure 3. At day 10, there was no significant difference between vascular and non-vascular staining for Col I and Sclerostin, whereas DMP-1 staining intensity was significantly lower in the vascular group compared to the non-vascular (Fig. 3 i, r, aa). By day 21, collagen I was significantly lower and staining intensity for sclerostin was significantly higher in the vascular group compared to the non-vascular (Fig. 3 i, aa). By day 21, there was a significant increase in the staining intensity of sclerostin in the vascular group compared to day 10 (Fig. 3 aa). In the vascular and non-vascular groups, von Kossa staining was higher at day 21 compared to day 10, and there were distinct mineralized nodules present only in the vascular groups (Fig. 3 bb-ii). Between day 10 and 21, there was a significant increase in the total calcium (normalized to weight of the construct) in the non-vascular group, but there was no significant difference in terms of calcium production between the vascular and non-vascular groups at day 10 or 21 (Fig. 3jj).

**Figure 3:**
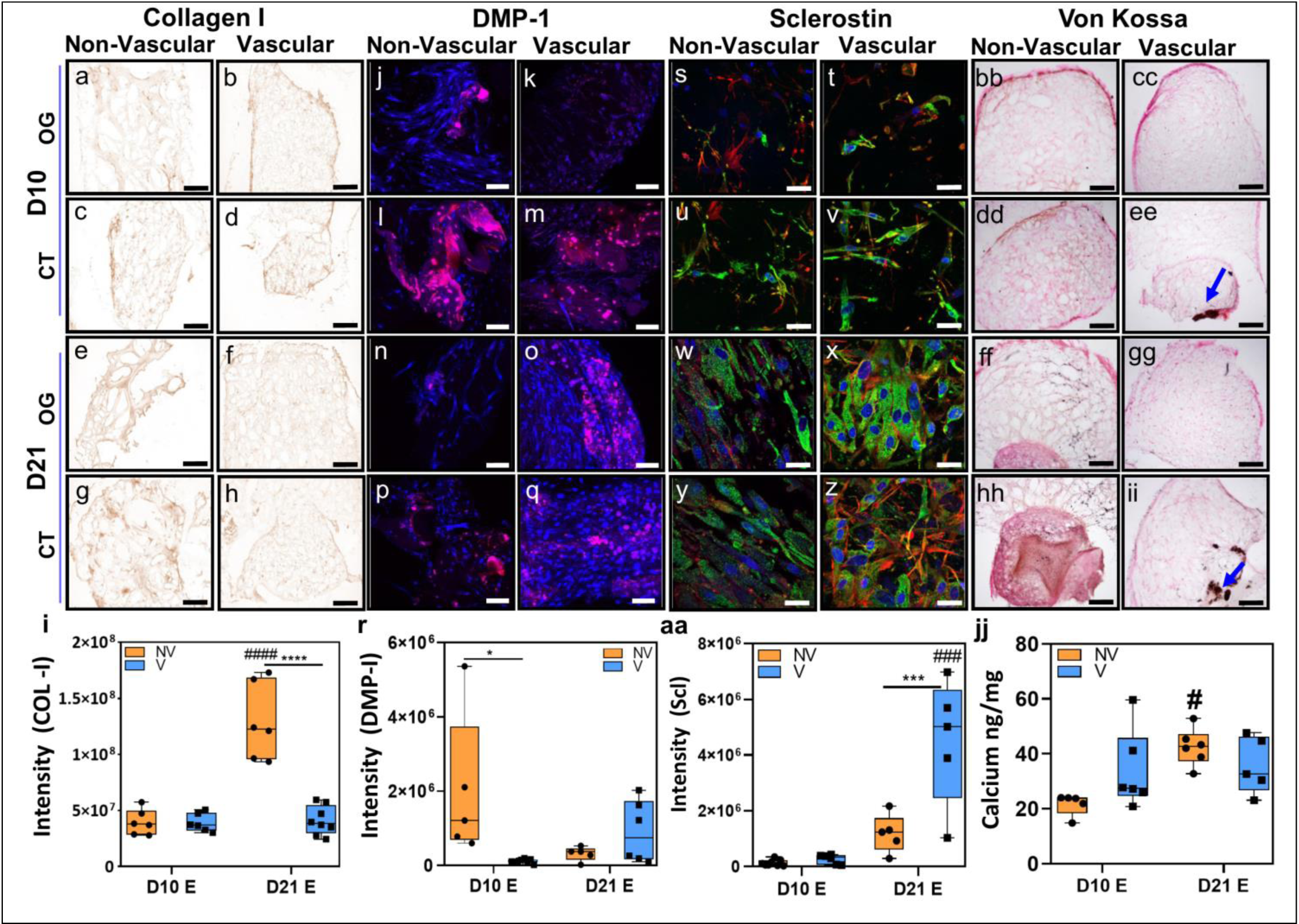
Vascular priming under estrogen supplementation promotes osteocyte differentiation and mineral production. Day 10 and 21 histology staining and quantification confirmed staining for (a-i) **Collagen I** (scale bar 500 µm), (j-r) **DMP-1** (Red), DAPI (Blue) (scale bar 100 µm), (s-aa) **Sclerostin** (green), actin (red) DAPI (blue) (scale bar 30 µm), (bb-ii) **Von Kossa** staining for mineral (scale bar 500 µm), (jj) **Total calcium** normalized to weight of construct. (OG: outer gel, CT: chondrogenic template). Significant differences indicated with * relative to non-vascular group and **#** relative to other time point. Two-way ANOVA, */# = p < 0.05, **/## = p < 0.01, ***/### = p < 0.001, ****/#### = p < 0.001.

We also reported that vascular priming under estrogen supplementation delays hypertrophy and apoptosis, when compared to the groups that were not cultured with vascular cells. Specifically, at day 10, there was significantly lower collagen X present in the vascular group compared to non-vascular (Fig. 4 i). For the vascular groups, collagen X intensity significantly increased between day 10 and 21 (Fig. 4 i). At day 21, there was a significant increase in BCL2 gene expression in the non-vascular group compared to day 10 (Fig. 4 k). BCL-2 was significantly lower in the vascular group compared to non-vascular at day 21 (Fig. 4 k). At day 21, the expression of caspase 3 significantly increased in the vascular group compared to day 10 (Fig. 4 m).

**Figure 4:**
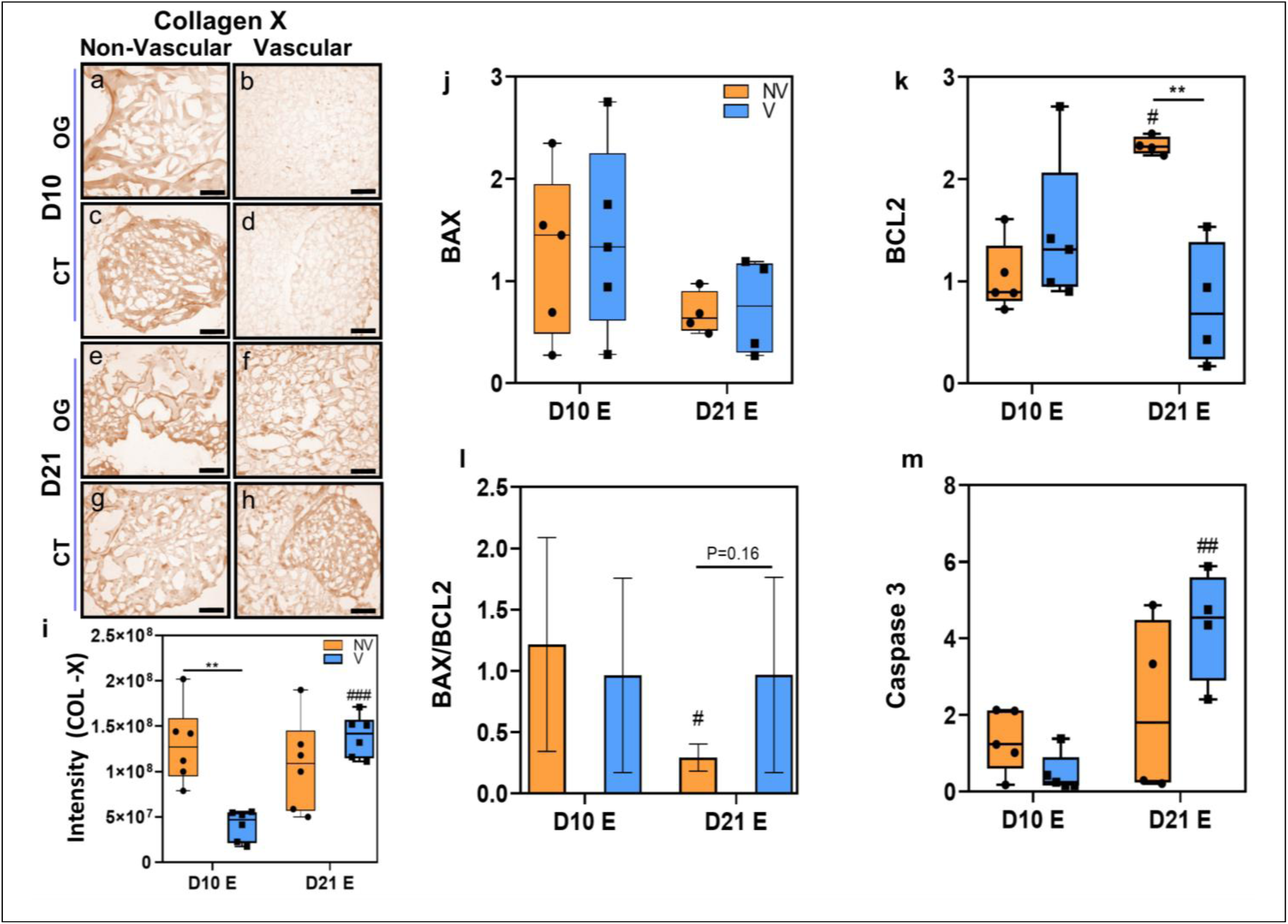
Vascular priming under estrogen supplementation delays hypertrophy (collagen X) and apoptosis. Day 10 and 21 histology staining and quantification for (a-i) **Collagen X** (scale bar 500 µm). (a-i) Gene expression for BAX, BCL2, BAX/BCL2, Caspase 3. Normalized to the non-vascular group at day 10. Significant differences indicated with * relative to non-vascular group and **#** relative to other time point. Two-way ANOVA, */# = p < 0.05, **/## = p < 0.01, ***/### = p < 0.001, ****/#### = p < 0.001.

### 3.3. Vascular priming under estrogen withdrawal leads to an increase in osteogenic differentiation and mineral production, which was associated with hypertrophy and apoptosis

Here we report that estrogen deficiency influences osteocyte differentiation and mineralization, with and without vascularization. At day 10 and 21, all the groups under estrogen withdrawal stained positive for collagen I, DMP-I, sclerostin and mineral. At Day 10 the intensity of collagen I and DMP-I staining was significantly lower in the vascular group compared to the non-vascular group (Fig. 5 i, r). By day 21, there was a significant increase in the staining intensity of collagen I in the vascular and non-vascular groups compared to day 10 (Fig. 5 i). At day 21 the staining intensity of collagen I was significantly lower in the vascular group compared to non-vascular group (Fig. 5 i). At day 21, the intensity of sclerostin staining significantly increased in the vascular and non-vascular group compared to day 10 (Fig. 5 aa). The Von Kossa staining showed intense mineral staining in the vascular group at day 21 (Fig. 5 jj), and the total calcium (normalized to weight of construct) was significantly higher than the non-vascular group (Fig. 5 jj).

**Figure 5:**
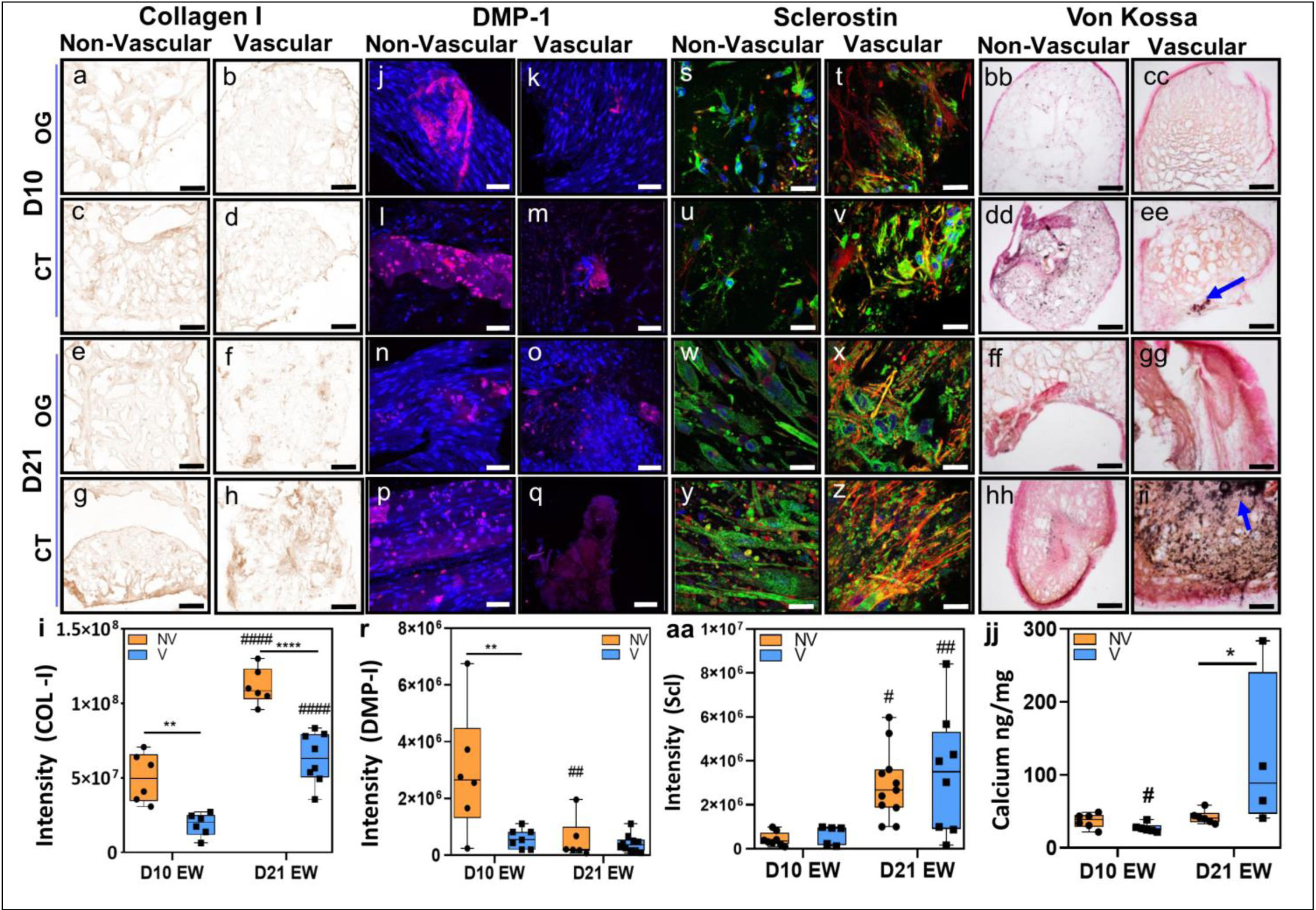
Vascular priming under estrogen withdrawal promotes osteocyte differentiation and significantly increases mineral deposition. Day 10 and 21 histology staining and quantification confirmed staining for (a-i) **Collagen I** (scale bar 500 µm), (j-r) **DMP-I** (Red), DAPI (Blue) (scale bar 100 µm), (s-aa) **Sclerostin** (green), actin (red) DAPI (blue) (scale bar 30 µm), (bb-ii) **Von Kossa** staining for mineral (scale bar 500 µm), (jj) **Total calcium** normalized to weight of construct. (OG: outer gel, CT: chondrogenic template) Significant differences indicated with * relative to non-vascular group and **#** relative to other time point. Two-way ANOVA, */# = p < 0.05, **/## = p < 0.01, ***/### = p < 0.001, ****/#### = p < 0.001.

These effects were accompanied by an increase in hypertrophy and apoptosis in the estrogen deficient and vascularized constructs. There was a significant increase in the intensity of collagen X in the vascular and non-vascular group at day 21 compared to day 10 (Fig. 6 i). At day 21, the gene expression of caspase 3 significantly increased in the vascular group compared to day 10 (Fig. 6 m). Notably, Caspase 3 expression was also significantly higher in the vascular group compared to non-vascular group at day 21 (Fig. 6 m).

**Figure 6:**
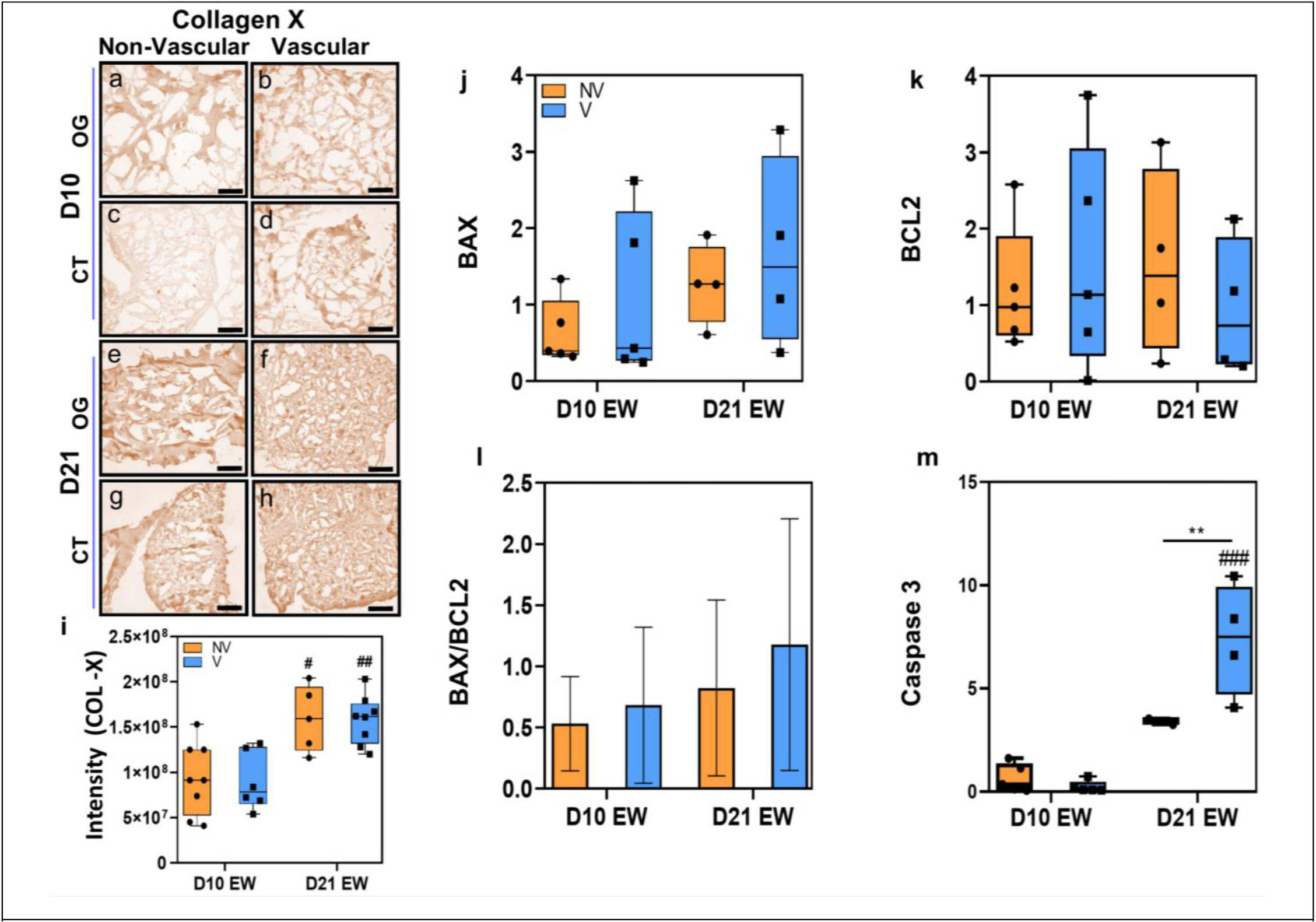
Vascular priming under estrogen withdrawal delays hypertrophy but significantly increases apoptosis. Day 10 and 21 histology staining and quantification for (a-i) Collagen X (scale bar 500 µm). (a-i) Gene expression for BAX, BCL2, BAX/BCL2, Caspase 3. Normalized to the non-vascular group at day 10. Significant differences indicated with * relative to non-vascular group and **#** relative to other time point. Two-way ANOVA, */# = p < 0.05, **/## = p < 0.01, ***/### = p < 0.001, ****/#### = p < 0.001.

### 3.4. Estrogen withdrawal alters vascular development

Here we report that estrogen withdrawal leads to changes in vascularisation, whereby mature vessel-like structures were only detected in the estrogen withdrawal group. By day 21, there was significant increase in the intensity of CD31 under estrogen supplementation compared to estrogen withdrawal (Fig. 7 a, c). There was no difference in the CD31 intensity in estrogen withdrawal group between day 10 and 21 (Fig 7 a, c). Similarly, estrogen deficiency significantly reduced the intensity of endomucin (EMCN) at day 10 (Fig 7 b, d). Confocal imaging showed that there were 3D multicellular vessel-like structures in the estrogen withdrawal group at day 21. These structures were random and rare in the vascular constructs, but were only present in the estrogen withdrawal group (Fig. 7 e, f, g, h). There was an increase in the gene expression of VEGF in the vascular estrogen withdrawal group at day 21 (Fig. 7 i).

**Figure 7:**
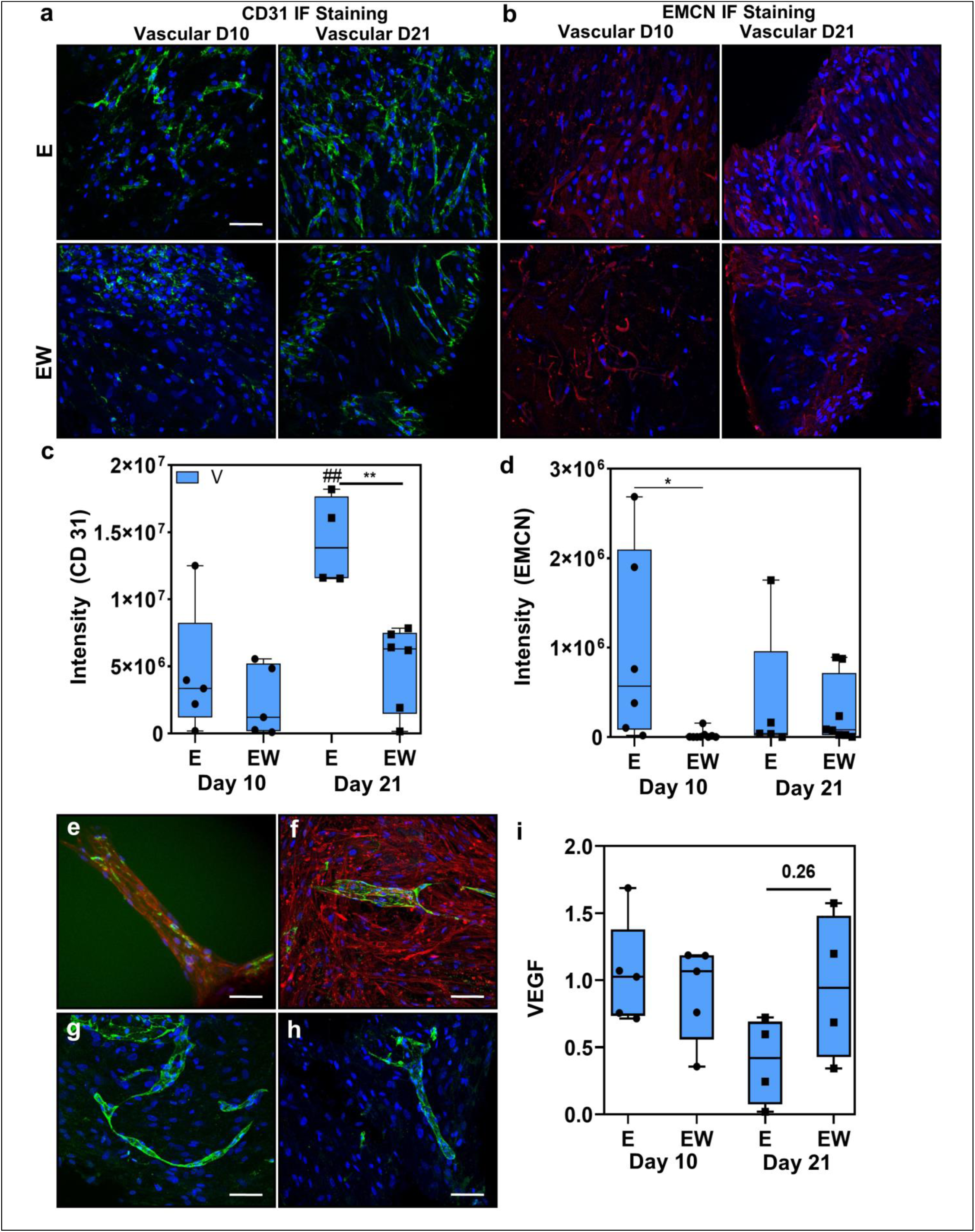
Distinct vessel-like structures (CD31+) are present in the estrogen withdrawal (postmenopausal) groups. **(a)** IFC staining for CD31 at day 10 and 21, CD31 (Green), DAPI (Blue) (scale bar 100μm) **(c)** Intensity of CD31. **(b)** IFC staining for EMCN (endomucin) at day 10 and 21, EMCN (Red), DAPI (Blue) (scale bar 100μm) **(d)** Intensity of EMCN. **(e, f, g, h)** 3D multicellular, vessel like structures in estrogen withdrawal group at day 21, CD31 (Green), DAPI (Blue), actin (Red). Gene expression for VEGF. Normalized to the vascular group estrogen supplemented group at day 10. Significant differences indicated with * relative to non-vascular group and **#** relative to other time point. Two-way ANOVA, */# = p < 0.05, **/## = p < 0.01, ***/### = p < 0.001, ****/#### = p < 0.001.

## 4. Discussion

The current study provides the first humanized bone mimetic model that successfully promoted vascularization and mineralization over an extended culture period, and recapitulated estrogen deficiency representative of the osteoporotic bone phenotype. We followed an endochondral ossification approach to induce formation of CD31+ 3D vessel-like structures that invaded the chondrogenic template. Our results demonstrate that vascular priming under estrogen supplementation enhances osteocyte differentiation and mineralization, while concurrently delaying hypertrophy and apoptosis. This suggests that estrogen plays a protective role in maintaining bone tissue homeostasis by regulating cellular maturation and promoting healthy mineral deposition. When our models were subjected to estrogen deficiency, we reported the development of distinct 3D vessel-like structures (CD31+) in the estrogen deficient postmenopausal and under these conditions vascularized bone demonstrated a significant increase in mineral deposition, which did not occur in non-vascularized bone or estrogen supplemented groups. Collagen X staining indicated that hypertrophic chondrocytes were responsible for endothelial cell invasion and vessel formation. This highlights a potential mechanism for heterogeneity in mineral distribution in osteoporotic bone. Collectively, these findings reveal the complex interplay between estrogen deficiency, vascularization, hypertrophy, and osteogenesis, offering new insights into bone health and disease progression under fluctuating estrogen levels.

This study has some limitations. Firstly, chondrogenesis and osteogenesis were encouraged through the inclusion of growth factors, which are not present during cartilage and bone formation during bone development *in vivo*. However, chondrogenic and osteogenic supplements have been widely used to encourage bone and cartilage regeneration by stem cells *in vitro* and *in vivo* studies [38–45]. Secondly, human umbilical cord derived endothelial cells from three different females were used to induce vascularization in our models, and so cannot represent a wider population. However, research shows that pooling of stem cells from different individuals does not affect the osteogenic and chondrogenic differentiation potential [46, 47]. Moreover, here we implemented an abrupt withdrawal of estrogen from our cultures that does not reflect the gradual decrease of estrogen in females during menopause [48]. However, ovariectomized animals also undergo a sudden depletion of circulating estrogen, but still are considered to be a good model of osteoporosis, thus our estrogen deficiency approach may considered equally applicable to study osteoporosis [49]. Finally, bone is a mechanically sensitive tissue, and although we showed the presence of sclerostin secreting, osteocyte-like mechanosensory cells in our model, we did not investigate the combined effect of mechanical stimulation and estrogen on our model. Future studies can apply our *in vitro* bone model to provide a mechanistic understanding of the role of mechanical stimulation during estrogen deficiency and osteoporosis. Nonetheless, our novel *in vitro* 3D humanized and vascularized bone model provides insights into the combined effects of estrogen withdrawal and vascularization on osteogenesis and mineralization.

Here we confirmed the successful development of a healthy (estrogen supplemented) vascularized and mineralized human bone model via endochondral ossification, which induced self-organization of vasculature, associated with hypertrophy (collagen X) and exacerbated mineralization. Previously, mesenchymal stem cells have been cultured in 3D systems to develop bone like phenotype to study osteogenesis [50, 51], bone metastases [52] and drug screening [53, 54], but no previous model mimicked the complex multicellular and vascularized micro-environment of bone. Our approach builds upon our previous studies [31, 32] by replicating endochondral ossification to develop a humanized chondrogenic template and induce vascularization and osteogenesis over long-term culture. In our healthy bone model, we confirm the presence of key osteogenic markers (collagens (II, X, I), DMP-1 and sclerostin), ALP activity and mineral production. During endochondral ossification, endothelial cells proliferate and invade the chondrogenic template under the influence of hypertrophic factors such as collagen X [29]. Similarly, in our healthy bone model, collagen X significantly increased over time, which coincided with the significant increase in the endothelial cell marker CD31. Previous studies have reported that endothelial cells support osteoprogenitors by releasing extracellular vesicles to promote their differentiation into osteoblasts [55–57]. Interestingly, our healthy vascular models showed significantly increased osteogenic differentiation (sclerostin staining) and similar mineral formation compared to non-vascular group. Although the non-vascular bone models had twice as many stem cells compared to vascular models, the osteogenic effect was pronounced in the vascular group, and so the stem cell population could not be independently driving this effect.

When the model was applied to study estrogen deficiency, we reported the development of distinct vessel-like structures (CD31+) in the postmenopausal 3D constructs. Moreover, during estrogen withdrawal vascularized bone demonstrated a significant increase in mineral deposition, which did not occur in non-vascularized bone. Endothelial cells have been previously reported to form tubular structures on 2D matrix and in 3D microfluidic systems [58, 59]. Here the formation of vascular structures was rare and random, but was only reported in the vascular estrogen withdrawal constructs. Estrogen plays important role in the vascular health [60] and estrogen supplementation can increase vascular proliferation [61]. In contrast to our findings, a recent *in vivo* study showed that estrogen deficiency, in an animal model decreases the number blood vessels and fatty acid metabolism, leading to increased number of adipocytes and decreased bone mineral [34]. This contrast might be explained by the differential response of human and mouse bone cells in the presence of vascular cells during estrogen deficiency *in vitro*. Moreover, our model did not include mechanical stimulation, whereas *in vivo* the animals maintained activity. Here we report a significant increase in the mineral deposition in the vascular and estrogen deficient group. We have previously established that estrogen deficiency can induce mineralization by murine osteoblasts [21, 62], causes apoptosis in murine osteocytes and also alters bone turnover and mineralization dynamics [18, 63, 64]. We also found a significant increase in the collagen X staining and caspase 3 expression in our vascular estrogen withdrawal model, which might explain the increased mineral production in that group. Previously, an *in vivo* study showed that ovariectomy induced estrogen deficiency causes alterations in the bone microarchitecture, and heterogeneous mineral distribution and composition [3, 65]. In our study, we report very high mineral deposition in the vascular and estrogen deficient model. This may explain the heterogeneity in bone mineralization after ovariectomy [3, 65], whereby estrogen deficiency leads to regions that become highly vascularized and form mineralized nodules, whereas in non-vascularized regions this effect is not observed. Thus, these findings reveal a potential mechanism for heterogeneity in mineral distribution in osteoporotic bone, whereby vascularized bone becomes highly mineralized whereas in non-vascularised regions this effect is not observed. Building on the findings of this study, future research should elucidate the specific molecular pathways that regulate altered calcium content and mineralization in bone tissue during estrogen deficiency. Understanding the precise role of vascular cells in these processes is particularly critical. To achieve this, advanced genomics and proteomics analyses will be essential for dissecting the contributions of each cell type within the complex, multicellular, and 3D microenvironment of bone. Additionally, integrating single-cell sequencing and spatial transcriptomics could provide deeper insights into cell-cell interactions and the dynamic changes that occur in response to estrogen withdrawal. These approaches will help to uncover novel therapeutic targets and strategies aimed at mitigating the effects of osteoporosis, particularly in postmenopausal women.

In conclusion, this study provides an advanced 3D vascularized and humanized *in vitro* bone model that shows the evidence of early stage osteogenesis and vascularization. When the model was applied to study estrogen deficiency, we reported the development of distinct vessel-like structures (CD31+) and significant mineral deposition in vascularized bone models. These findings reveal a potential mechanism for heterogeneity in mineral distribution in osteoporotic bone, whereby vascularized bone becomes highly mineralized but non-vascularised bone does not. Overall, this approach will inform the development self-organizing *in vitro* bone models that may serve as surrogate for drug testing and precision medicine.

## Acknowledgments

This research was funded by European Research Council (ERC) Consolidator grant (MEMETic 863795). The authors acknowledge the facilities and technical assistance of the Centre for Microscopy and Imaging at the University of Galway (https://imaging.universityofgalway.ie). The authors acknowledge Dr. Colin Murphy for providing bone marrow specimen under Ethical under ethical approval and informed consent to the patients (Research Ethics Committee, University of Galway). The authors would also like to acknowledge Smart Servier Medical Art (https://smart.servier.com/) for their image bank used to produce figure 6.

## Data availability

Raw data can be made available upon responsible request.

## Author contributions

Project administration and funding acquisition, L.McN; Conceptualization, B.M.M.M., S.M.N., L.McN.; methodology, B.M.M.M., M.K., S.M.N.; formal analysis and data interpretation, B.M.M.M., M.K, S.M.N., L.McN.; writing, review and editing B.M.M.M., L.McN.; final proof reading and approval B.M.M.M., M.K, S.M.N., L.McN; L.McN; All authors have read and agreed to the published version of the manuscript.

---

**Supplementary Figure 1.**
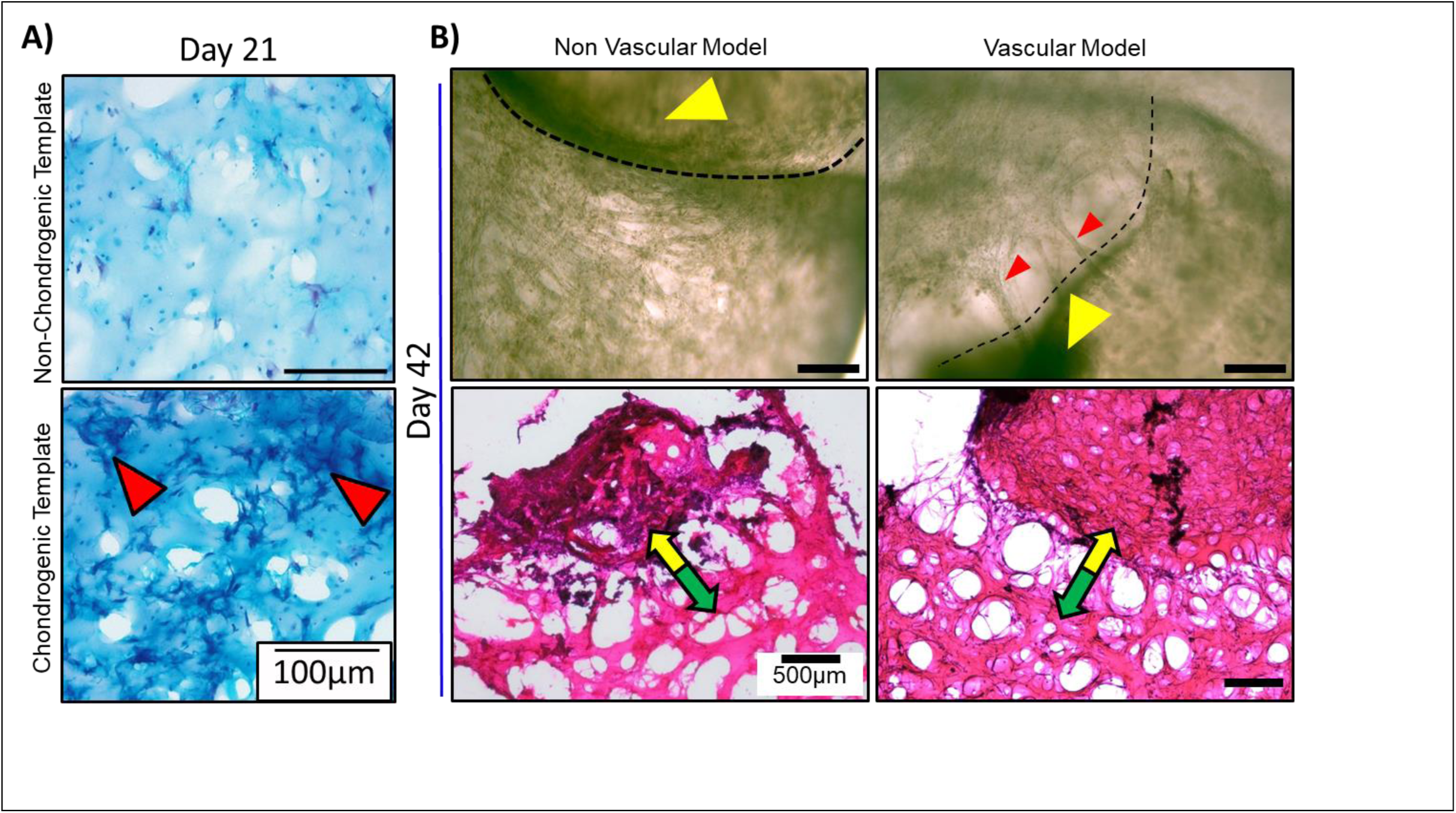
A) Alcian blue staining for chondrogenesis B) Phase contrast microscopy and H&E staining.

**Supplementary Figure 2.**
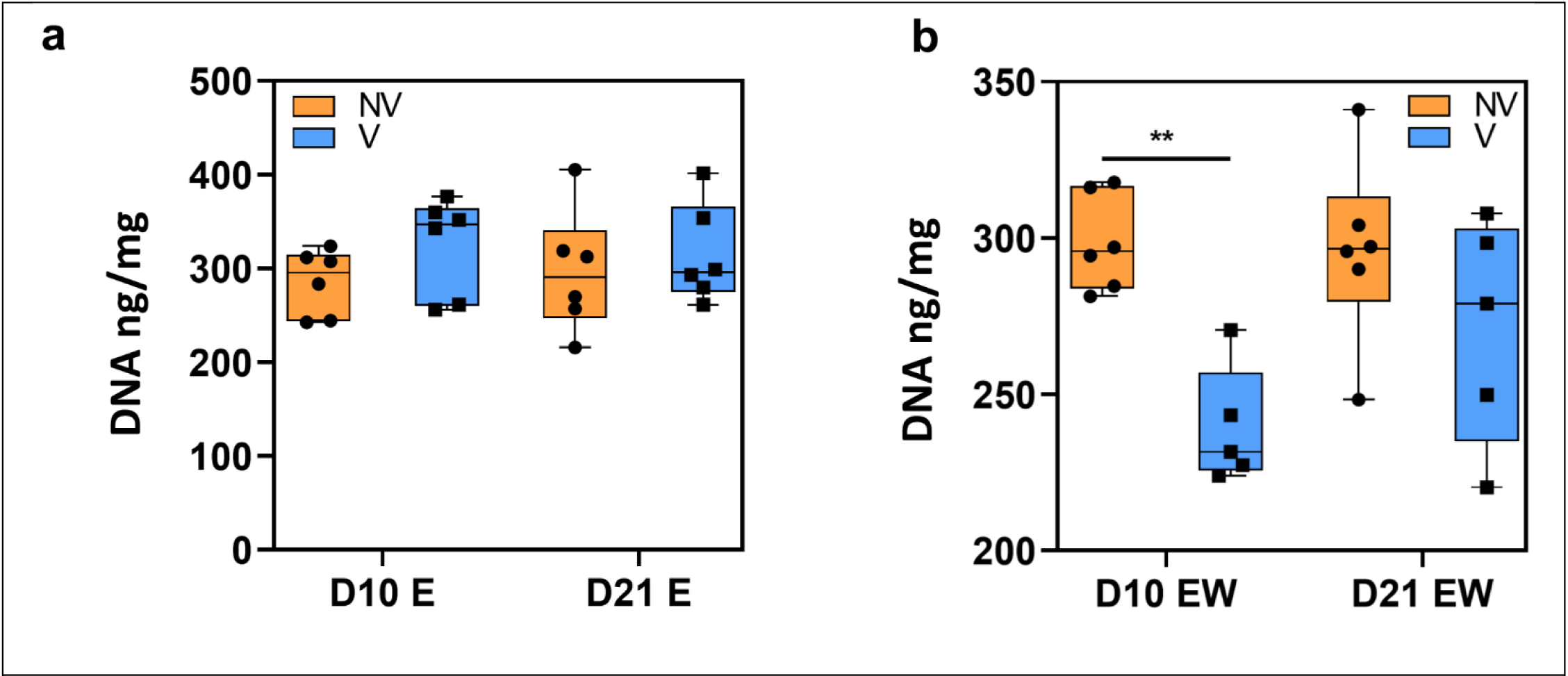
Total DNA under (a) estrogen supplementation and (b) estrogen withdrawal. Significant differences indicated with * relative to non-vascular group and # relative to other time point. Two-way ANOVA, */# = p < 0.05, **/## = p < 0.01, ***/### = p < 0.001, ****/#### = p < 0.001.

**Supplementary Figure 3.**
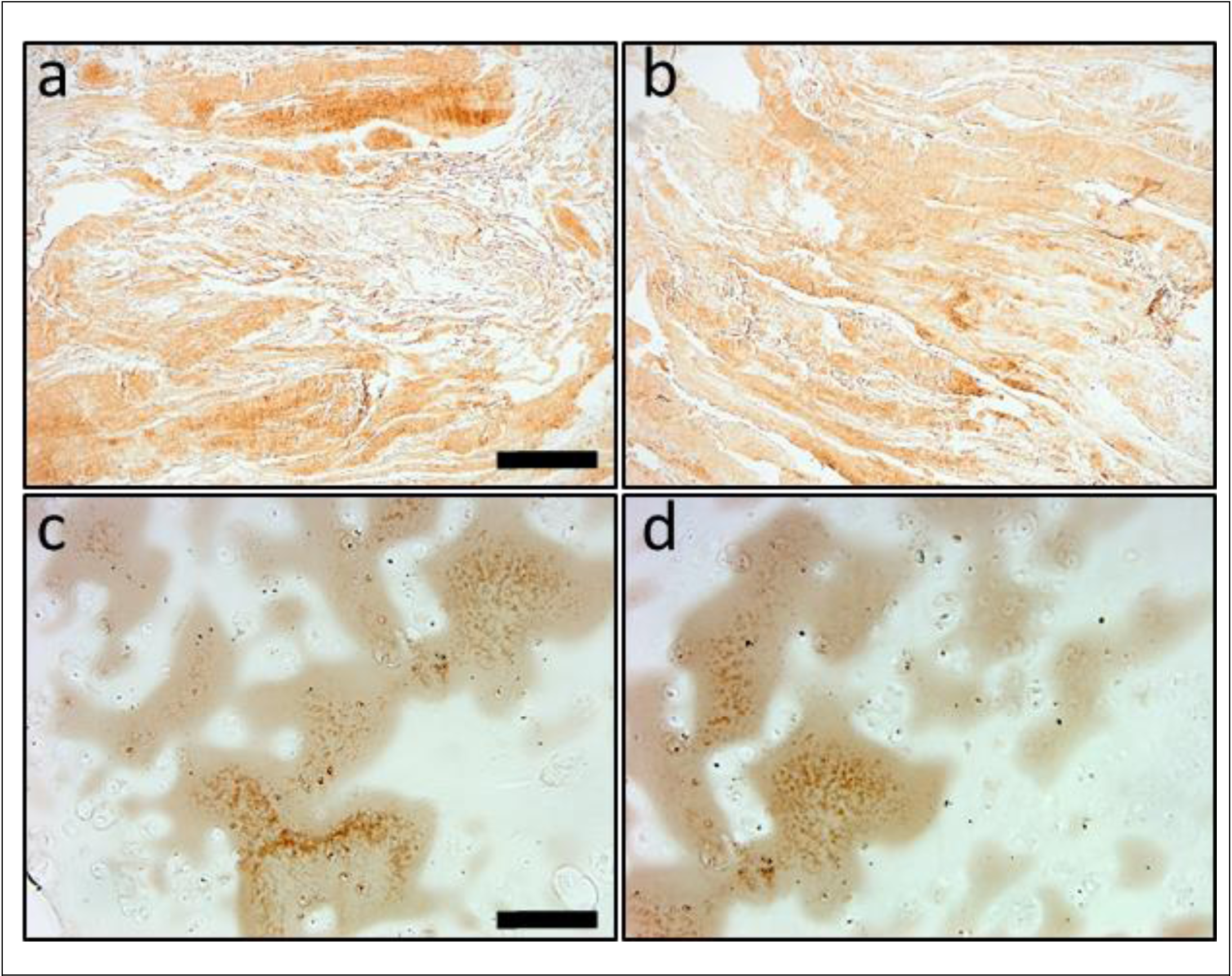
(a, b) Immunohistochemistry staining of human ligament sections for Collagen I (scale bar 500µm). (c, d) Immunohistochemistry staining of human cartilage sections for Collagen X (scale bar 500µm).

